# ^225^Ac-labeled CD33-targeting antibody reverses resistance to Bcl-2 inhibitor venetoclax in acute myeloid leukemia models

**DOI:** 10.1101/2020.10.09.334151

**Authors:** Ravendra Garg, Kevin J.H. Allen, Wojciech Dawicki, Eileen M. Geoghegan, Dale L. Ludwig, Ekaterina Dadachova

## Abstract

**Purpose:** Despite the availability of new drugs, many patients with acute myeloid leukemia (AML) do not achieve remission and outcomes remain poor. Venetoclax is a promising new therapy approved for use in combination with a hypomethylating agent or with low dose cytarabine for the treatment of newly diagnosed older AML patients or those ineligible for intensive chemotherapy. ^225^Actinium-lintuzumab (^225^Ac-lintuzumab) is a clinical stage radioimmunotherapy targeting CD33 that has shown evidence of single agent activity in relapsed/refractory AML. Increased expression of MCL-1 is a mediator of resistance to venetoclax in cancer.

**Experimental design:** Here we investigated the potential for ^225^Ac-lintuzumab-directed DNA damage to suppress MCL-1 levels as a possible mechanism of reversing resistance to venetoclax in two preclinical in vivo models of AML.

**Results:** We demonstrated that ^225^Ac-lintuzumab in combination with venetoclax induced a synergistic increase in tumor cell killing compared to treatment with either drug alone in venetoclax-resistant AML cell lines through both an induction of double-stranded DNA breaks (DSBs) and depletion of MCL-1 protein levels. Further, this combination led to significant tumor growth control and prolonged survival benefit in venetoclax-resistant in vivo AML models.

**Conclusions:** There results suggest that the combination of ^225^Ac-lintuzumab with venetoclax may be a promising therapeutic strategy for the treatment of patients with venetoclax-resistant AML.

## INTRODUCTION

Although standard induction therapy with cytarabine and an anthracycline produces complete remissions (CR) in 60% to 80% of younger adults with acute myeloid leukemia (AML), long-term survival is seen in only 25% to 50% of patients ^1^. Following relapse, salvage chemotherapy produces remissions in only 20% to 25% of patients. Further, while allogeneic hematopoietic cell transplantation (HCT) can result in long-term survival in approximately 30% to 40% of patients with relapsed AML, many patients are not suitable candidates for transplant due to age, comorbidities, or lack of a matched donor ^1^. The prognosis for older patients is even worse, with a 5-year median survival rate of 5% for patients older than age 65 ^2^. Within the last few years, following decades of limited advancement in the treatment of AML, several new targeted therapies have been approved in the US ^3,4^. Notably, the BCL2 inhibitor venetoclax (ABT-199), was approved for use in combination with a hypomethylating agent (HMA) or with low-dose cytarabine (LDAC) for the treatment of newly diagnosed patients with AML who are age 75 years or older or are ineligible for intensive chemotherapy ^5,6^. Approval was based on the M14-358 and M14-387 phase Ib/II trials. In M14-358 the combination of venetoclax with azacitidine led to a complete remission (CR) rate of 37% and a CR with partial hematological recovery (CRh) rate of 24% ^5^. For the combination of venetoclax and decitabine, the rates were 54% and 7.7%, respectively. In the M14-387 trial, venetoclax in combination with low-dose cytarabine (LDAC) led to 21% CR and CRh rates ^6^. For patients receiving all dosages of venetoclax in these studies, median overall survival was 17.5 months ^5^. Despite the significant benefit to patients in this population, not all patients respond to initial therapy with venetoclax and most patients will eventually progress. Preclinical studies have investigated mechanisms of resistance, which are supported by clinical results in patients with progressive disease. Importantly, venetoclax as a B cell lymphoma 2 (BCL-2) selective inhibitor, does not inhibit other BCL family members such as myeloid cell leukemia 1 (MCL-1) or B lymphoma extra-large (BCL-XL). Mechanistically, overexpression of these other anti-apoptotic BCL-2 family members, in particular MCL-1, or their up-regulation in response to venetoclax have been shown to mediate resistance to venetoclax in leukemia, lymphoma and multiple myeloma ^7-11^. Further, MCL-1 has been shown to be upregulated in AML patients at relapse following induction chemotherapy ^12^. Strategies to reduce MCL-1 levels may therefore dramatically prolong the response to venetoclax and re-sensitize resistant tumors to venetoclax therapy. MCL-1 protein has a very short half-life of less than 1 hour and is therefore sensitive to changes in RNA or protein synthesis ^13^. To that end, genotoxic stress as a result of DNA damage, e.g. by UV or ionizing radiation, or chemo-induced, can effect a reduction in MCL-1 levels via inhibition of protein synthesis ^14,15^. In turn, the combination with chemotherapy or ionizing radiation can increase sensitivity to BCL-2 inhibitors in preclinical tumor cell lines and patient samples by a reduction in MCL-1 levels ^7,14,16^. ^225^Ac-lintuzumab is a clinical stage radioimmunotherapy targeting CD33 that has shown evidence of single agent activity in relapsed/refractory AML ^17-19^ and is being actively investigated in combination clinical studies in the treatment of relapsed or refractory AML. ^225^Ac-lintuzumab delivers the ^225^Ac payload, a high energy, short path length, alpha emitting radionuclide directly to CD33 positive myeloid tumor cells creating lethal double strand breaks in DNA and leading to selective tumor cell killing ^17^. As little as on alpha hit from ^225^Ac radionuclide can potentially kill a tumor cell, and the short path length focuses its radiation energy on targeted tumor cells, limiting exposure and damage to adjacent normal tissue ^20^. While external beam radiation has been combined with venetoclax in preclinical models, there is the potential to damage normal tissue. Further, scheduling radiation treatment relative to venetoclax administration may also be a challenge. As a potent inducer of DNA damage in targeted tumor cells, it is anticipated that ^225^Ac-lintuzumab may mediate effective down-modulation of MCL-1 leading to a sensitization of AML cells to venetoclax irrespective of inherent resistance to the BCL-2 inhibitor. Thus, the combination of potent targeted alpha radioimmunotherapy (RIT) plus venetoclax may be an effective combination strategy in in AML. Here we describe the results of our *in vitro* and *in vivo* studies which evaluated ^225^Ac-lintuzumab plus venetoclax in combination in established AML tumor cell lines exhibiting varying sensitivity to venetoclax. Unlike other strategies to deplete MCL-1 in combination with venetoclax, our results demonstrate that ^225^Ac-lintuzumab targeted internal alpha radiation exerts a dual mechanism of action, effecting potent single agent tumor killing through DNA double strand breaks, and the reduction in anti-apoptotic proteins such as MCL-1, leading to re-sensitization of tumor cells to venetoclax and potent anti-tumor activity in preclinical models.

## MATERIALS AND METHODS

### Antibodies, reagents, and cell lines

Humanized anti-CD33 antibody, lintuzumab was provided by Actinium Pharmaceuticals, Inc. (New York, NY, USA) ^17^. Venetoclax was purchased from Selleckchem (Huston, TX, USA). For animal studies venetoclax was formulated in 60% Phosal 50PG (Lipoid), 30% polyethylene glycol 400 (Sigma), 10% ethanol. ^225^Ac in anhydrous nitrate form was procured from Oak Ridge National Laboratory, USA. Bifunctional chelating agent p-SCN-Bn-DOTA (DOTA) was purchased from Macrocyclics (Plano, TX, USA). AML cell lines MOLM-13, and OCI-AML3 were purchased from Deutsche Sammelung von Mikroorganismen und Zellkulturen (DSMZ, Braunschweig, Germany) and U937 from the ATCC (Manassas, VA, USA). All cell lines were cultured in RPMI (Thermo Fisher Scientific, Waltham, MA, USA) supplemented with 10% fetal bovine serum (Sigma-Aldrich, St. Louis, MO, USA) and 1% antibiotic/antimycotic (Thermo Fisher Scientific). Cells were kept at 37°C in a 5% CO_2_ incubator. CD33 expression by the AML cell lines was determined by flow cytometry. Cells were stained with PE-conjugated anti-human CD33 or isotype control mAbs (BD Pharrmingen, San Jose, CA).

### Lintuzumab antibody conjugation and radiolabeling with ^225^Ac

The antibody lintuzumab was conjugated to the bifunctional chelator *p*-SCN-Bn-DOTA (Macrocyclics) by methods described previously with some modifications ^21^. In brief, 500 ug of lintuzumab was conjugated to the linker-chelator using a 50-fold molar excess of *p*-SCN-Bn-DOTA and incubated at 37°C for 1.5 h in sodium carbonate buffer. Upon completion of the reaction the lintuzumab-DOTA conjugate was then exchanged into the 0.15 M ammonium acetate buffer at 4°C. The molar ratio of chelator to antibody in DOTA-lintizumab was determined to be approximately 10:1 by MALDI-TOF (University of Alberta, Canada). The retention of immunoreactivity of DOTA-lintuzumab conjugate was confirmed by CD33 binding ELISA.

For radiolabeling, ^225^Ac nitrate was taken up in diluted HCl and incubated with lintuzumab-DOTA conjugate for 60 min at 37°C to achieve a 1:1 µCi/µg specific activity. The reaction was quenched by the addition of 0.05 M DTPA solution and the percentage of radiolabeling yield measured by instant thin layer chromatography (iTLC) SG-Glass microfiber chromatography paper impregnated with Silica Gel (Agilent Technologies, CA, USA, Cat. No.: SG10001) 24 h post-labeling when secular equilibrium is reached between ^225^Ac and its daughter isotopes using a 2470 Wizard2 Gamma counter (Perkin Elmer, MA, USA) calibrated for the ^225^Ac and ^213^Bi emission spectra. Radiolabeling yields were greater than 99% and no further purification was required. The R_F_ of the main product is 0 whereas the R_F_ of the ^225^Ac-DTPA is 1. Any free ^225^Ac-will be bound by the DTPA used during the quench due to the molar excess of DTPA added. The iTLC is read 24 h after elution to allow for ^225^Ac and its daughters to reach secular equilibrium where a true ratio of antibody bound ^225^Ac and “free” ^225^Ac can be measured. The stability of ^225^Ac-lintuzumab (formally known as ^225^Ac-HuM195) in vitro and in vivo was determined previously ^22^ and showed no leakage of Ac from the product.

### In vitro AML cell cytotoxicity assay

U937 and OCI-AML3 cells were cultured with 0-10,000 nM doses of venetoclax, whereas, MOLM-13 cells were cultured with doses ranging from 0-400 nM. AML cells were individually treated with unlabeled lintuzumab (0.1 μg, 0.74 KBq /0.02 μg, 1.48 KBq/0.04 μg, 2.22 KBq/0.06 μg and 3.7 KBq/0.1 μg ^225^Ac-lintuzumab for 1 h, washed, and cultured. For combination experiments, AML cell lines were treated with 1.48 KBq ^225^Ac-lintuzumab for 1 h, washed, and then incubated with 50 nM (MOLM-13) or 500 nM (U937 and OCI-AML3) venetoclax. Cells were cultured for 72 h and then cell death was measured using the tetrazolium dye (2,3)-bis-(2-methoxy-4-nitro-5-sulphenyl)-(2H)-terazolium-5-carboxanilide (XTT) assay (Sigma) according to manufacturer’s instruction. After 72 h treatment, cells were incubated in the presence of 50 μl XTT (1 mg/ml), and 4 μl menadione (1 mM) for another 3 h and the absorbance was read at 492 nm. The XTT proliferation assay was selected for use, as it has been widely utilized in cancer drug development studies to determine their cytotoxic effects. ^23,24^

### Phospho-H2A.X assay for detection of DNA double strand breaks

The presence of phosphorylated H2A.X was measured by flow cytometry according to the manufacturer’s instructions (Upstate Cell Signaling, Millipore Sigma). Briefly, fixed and permeabilized cells were stained with FITC conjugated anti-phospho-H2A.X (Ser139) or isotype control mouse IgG. Stained samples were analyzed with a CytoFLEX flow cytometer (Beckman Coulter, Mississauga, ON) and data were analyzed using FlowJo software (Tree Star Inc., Ashland, OR).

### Quantitative Western Blot

Cells were lysed in ice-cold lysis buffer (Cell Signaling Technology, Denver, MA, USA) containing protease inhibitor PMSF (Cell Signaling Technology). Total protein was extracted in sample buffer (Cell Signaling Technology), sonicated, and boiled at 95°C for 10 min. Equal amounts of protein were loaded to 10% Sodium Dodecyl Sulphate Polyacrylamide gel and Western blot analysis was performed according to standard protocol with antibodies against human MCL-1, BCL-2, BCL-XL, and anti-mouse α-tubulin (Cell Signaling Technology). Secondary detection was performed using LI-COR IRDye antibodies (IRDye 680-labeled anti-rabbit and IRDye 800-labeled anti mouse), visualized by the LI-COR Odyssey Imaging System (Lincoln, NE, USA) and quantified by Image J Studio lite software (LI-COR Biosciences).

### In vivo efficacy studies

Animal studies were approved by the University of Saskatchewan’s Animal Research Ethics Board and followed to the Canadian Council on Animal Care guidelines for humane animal use. Tumor xenografts were established in female SCID mice (CB17/Icr-Prkdc^scid^/IcrIcoCrl, obtained from Charles River Laboratories, Saint-Constant, QC, Canada) by subcutaneous injection of 2×10^6^ OCI-AML3 or U937 cells into the right flank. Tumor growth was measured with electronic calipers every 3 days (volume=length × width^2^/2). When tumors reached an average volume of ∼200 mm^3^, tumor-bearing mice were randomized into five treatment groups of 5 animals and were treated with on Day 1: IP injection of 0.4 μg unlabeled lintuzumab; IP injection of 7.4 KBq ^225^Ac-lintuzumab; venetoclax (200 mg/kg) by oral gavage; combination of venetoclax and 7.4 KBq ^225^Ac-lintuzumab; or left untreated. Mice in venetoclax alone and in combination treatment groups continued to receive venetoclax via oral gavage once daily over a period of 20 days. Mice body weight and tumor volume were recorded three times per week. The animals were humanly euthanized when they experienced excessive weight loss, or any tumor reached 4,000 mm^3^ volume or became necrotic. For safety study, blood was collected from euthanized mice and analyzed for liver toxicity (aspartate aminotransferase; AST and alanine transaminase; ALT) and kidney toxicity (creatinine and blood urea nitrogen; BUN).

### Statistical analysis

GraphPad Prism 7 was used to analyze all the data (GraphPad Software, Inc., La Jolla, CA, USA). Differences among the groups were assessed using Student t-tests and one-way ANOVAs for multiple comparisons. Differences were considered significant at P<0.05

## RESULTS

### Single agent cytotoxicity of venetoclax and ^225^Ac-lintuzumab in AML cell lines

Initially, two venetoclax resistant (OCI-AML3 and U937) and one sensitive (MOLM-13) AML cell lines were screened for CD33 expression. Flow cytometry analysis revealed high expression of CD33 in all three AML cell lines (Supplementary Fig. S1). To determine the optimal dose for in vitro combination studies, we evaluated the sensitivity of these cell lines to single-agent venetoclax or ^225^Ac-lintuzumab. The AML cell lines were treated individually with increasing concentrations of venetoclax or ^225^Ac-lintuzumab and cell viability measured after 72 h by XTT assay. U937 and OCI-AML3 cells began to show evidence of cytotoxicity at 500 nM venetoclax treatment, whereas, MOLM-13 cells were shown to be sensitive to concentrations as low as 5nM (Fig. 1A). The IC_50_ for venetoclax in MOLM-13 was determined to be 8.5 nM in comparison to venetoclax resistant lines U937 (IC_50s_ 663 nM) and OCI-AML3 (IC_50s_ 1115 nM). Treatment of U937 and OCI-AML3 cells with ^225^Ac-lintuzumab showed modest sensitivity following 1-hour exposure to the antibody radio-conjugate. MOLM-13 was shown to be more sensitive to ^225^Ac-lintuzumab at concentrations as low as 1.48 KBq (Fig. 1B).

**Figure 1:**
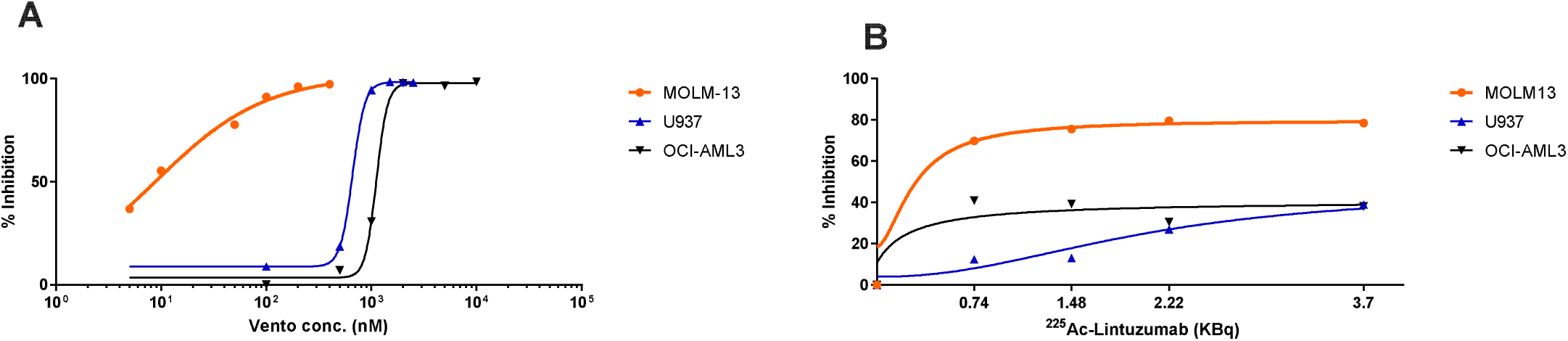
Single agent cellular cytotoxicity of venetoclax or ^225^Ac-Lintuzumab treatment in AML cell lines. The AML cell lines (MOLM-13, U937 and OCI-AML3) were treated individually with increasing dose of venetoclax (A) or ^225^Ac-Lintuzumab (B) and cell viability was measured using the XTT assay after 72h post-treatment. For ^225^Ac-Lintuzumab treatment, cells were treated with serial dilutions of ^225^Ac-Lintuzumab for 1 h, washed, and then cultured for 72 h before performing the XTT assay. Data are shown as percentage of inhibition of three replicates per cell type.

### ^225^Ac-lintuzumab and venetoclax combination shows synergetic cytotoxicity towards both sensitive and resistant AML cell lines

Based on results of single agent cytotoxicity of venetoclax and ^225^Ac-lintuzumab, we performed studies to assess the potential for enhanced cell killing with the combination of ^225^Ac-lintuzumab with venetoclax in both sensitive and resistant AML lines. In vitro combination cell cytotoxicity studies utilized IC_50_ concentrations determined from the single agent analyses: venetoclax, 50 nM (MOLM-13) or 500nM (U937 and OCI-AML3) and 1.48 KBq (^225^Ac -lintuzumab). The cells were first exposed to ^225^Ac-lintuzumab in the presence of venetoclax, then washed and subsequently incubated in medium containing fresh venetoclax. Inhibition of cell growth was measured after 72 h post-treatment by XTT assay. Treatment with ^225^Ac-lintuzumab combined with venetoclax resulted in significantly greater tumor cell killing as compared to individual treatment in all three cell lines (Fig 2). Importantly, the significant enhancement of cell killing in the two venetoclax resistant lines suggests that the combination of venetoclax with ^225^Ac-lintuzumab may affect re-sensitization of these lines to venetoclax.

**Figure 2.**
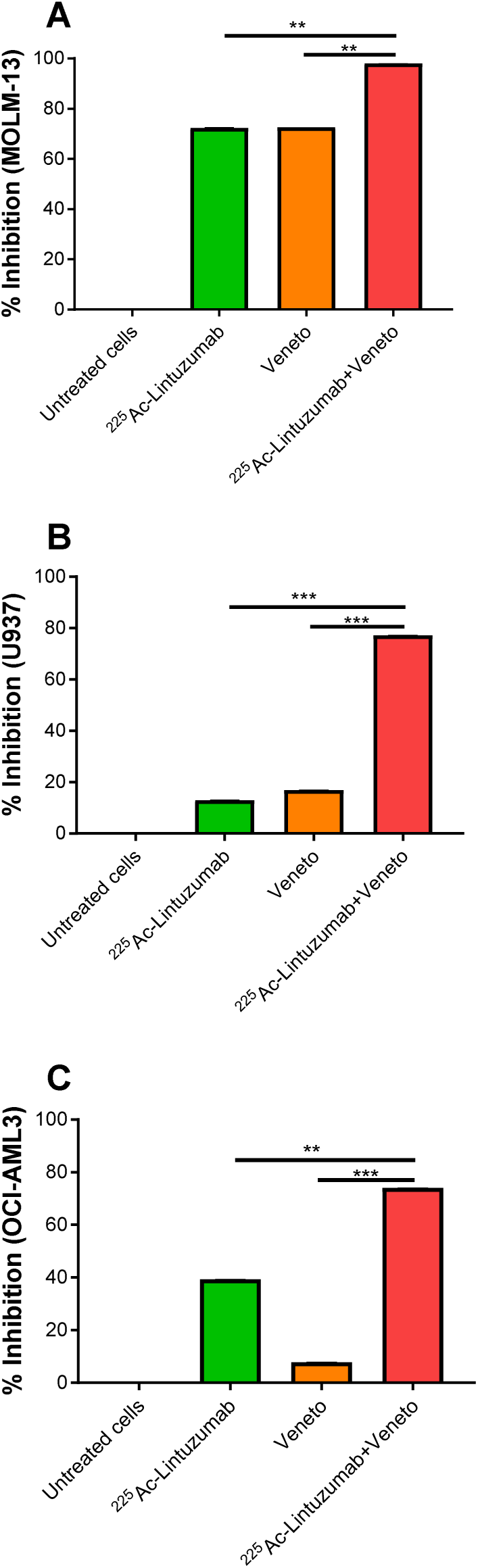
Combination treatment with ^225^Ac-Lintuzumab and venetoclax (veneto) induces enhanced cytotoxicity in AML cell lines. MOLM-13 (A), U937 (B) and OCI-AML3 (C) cell lines were pre-treated with ^225^Ac-Lintuzumab for 1 h, washed, and then incubated with 50 nM (MOLM-13) or 500nM (U937 and OCI-AML3) venetoclax for 72 h. Cell viability was measured by XTT assay. Data are shown as percentage of inhibition of three replicates per cell type. **p<0.01, *** p<0.001.

### Initiation of double strand breaks in AML cells exposed to ^225^Ac-lintuzumab

The ^225^Ac-lintuzumab carries the ^225^Ac radionuclide generating potent high linear energy transfer alpha particles which can cause lethal DNA double strand breaks (DBSs). Upon induction of DNA damage, the nucleosomal histone protein H2A.X is rapidly phosphorylated at serine 139 to *γ*-H2A.X at the DSBs site, making phosphorylated *γ*-H2A.X a sensitive marker for detection of DNA DSBs ^25^. To define the potential mechanisms by which ^225^Ac-lintuzumab induces significantly enhanced cellular cytotoxicity in AML cell lines, we screened for phosphorylated H2A.X following exposure to the antibody radio-conjugate. Treatment of both U937 and OCI-AML3 cells with ^225^Ac-lintuzumab alone or its combination with venetoclax induced high levels of *γ*-H2A.X as compared to venetoclax with synergistic effect observed for combination treatment (Fig. 3).

**Figure 3.**
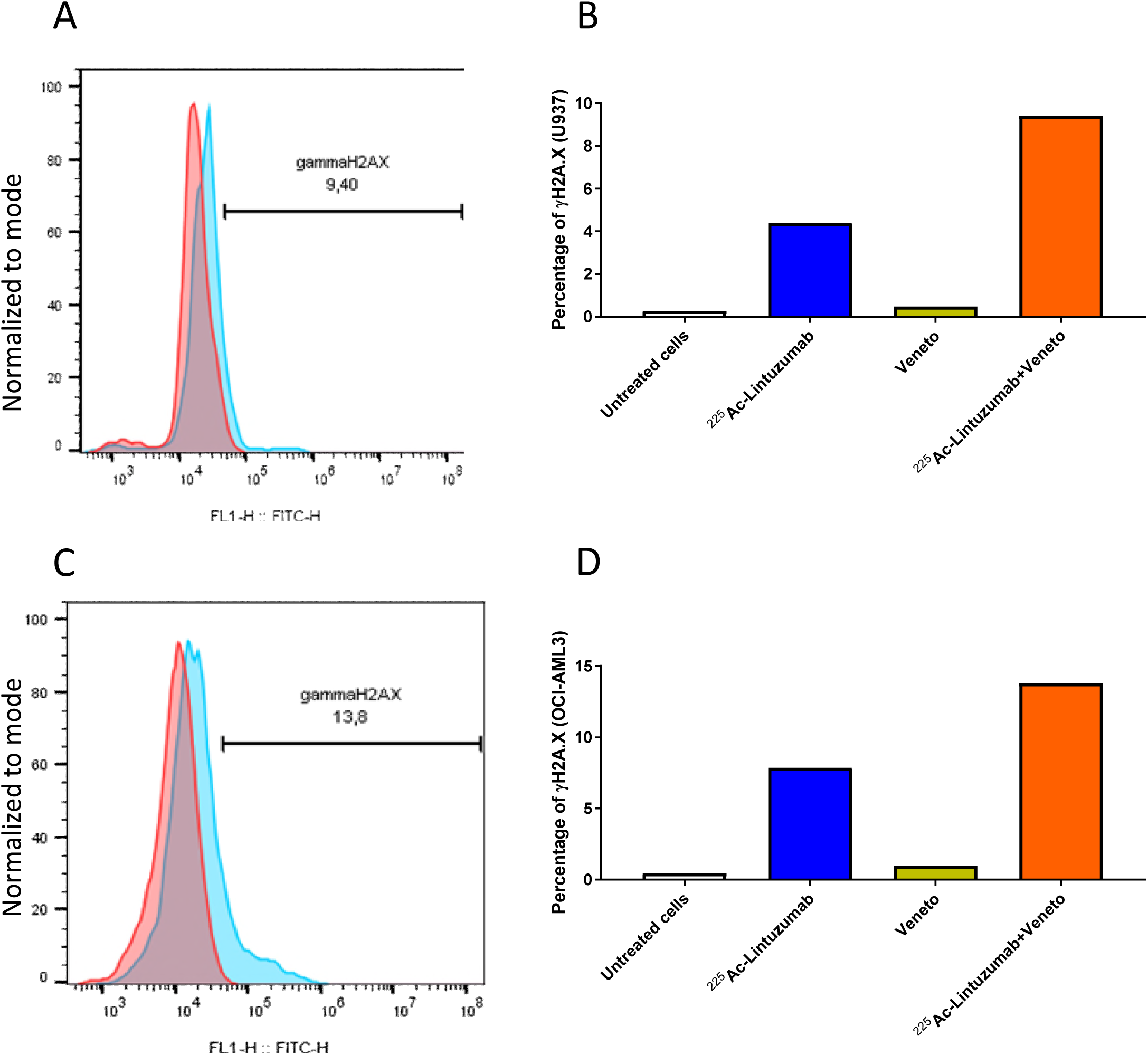
^225^Ac-Lintuzumab induces double-stranded DNA breaks in AML cell lines. U937 (A and B) and OCI-AML3 (C and D) cells were treated with media, venetoclax alone (500 nM), ^225^Ac-Lintuzumab (1.48 KBq) or combination with venetoclax. The presence of phosphorylated gamma H2A.X was measured by flow cytometry after 72 h. On the histogram plots (A and C), red line shows binding of isotype control antibody and blue line shows binding of γH2A.X antibody.

### ^225^Ac-lintuzumab reduces MCL-1, BCL-2 and BCL-XL levels in treated AML cells

It has been reported that genotoxic stress may inhibit the expression of BCL-2 family proteins such as MCL-1 ^14,15^, a protein implicated in resistance to BCL-2 inhibitors such as venetoclax. To evaluate the potential for ^225^Ac-lintuzumab to modulate MCL-1 levels, we assessed the levels of anti-apoptotic proteins in OCI-AML3 cells via western blot. We determined that transient exposure of OCI-AML3 cells to ^225^Ac-lintuzumab significantly reduced the levels of not only MCL-1, but also BCL-XL, and to a lesser extent BCL-2, in a dose dependent manner. This decrease in BCL-2 family protein levels may serve as a mechanism by which this AML cell line can become re-sensitized to venetoclax following combination treatment with ^225^Ac-lintuzumab (Fig. 4).

**Figure 4:**
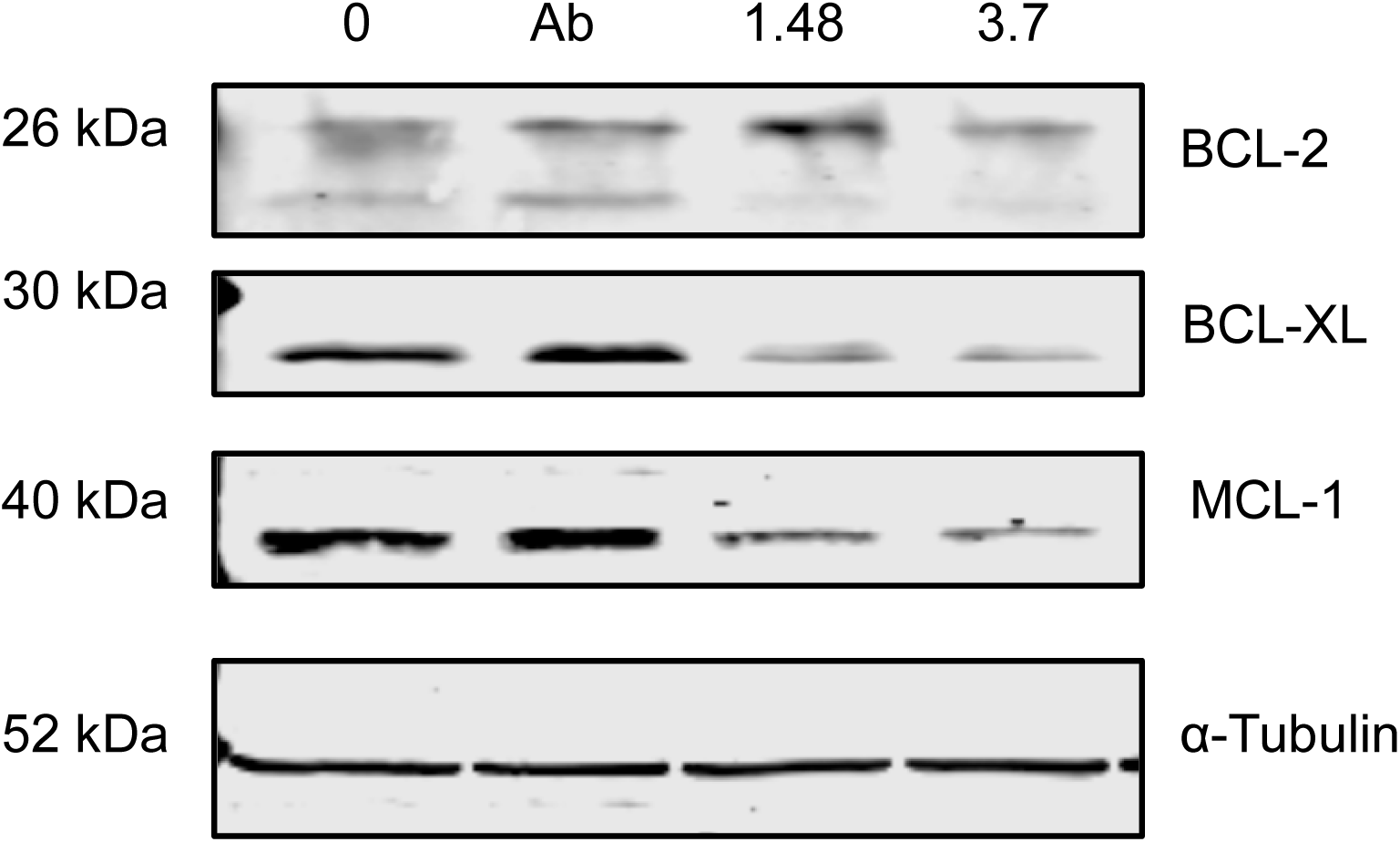
^225^Ac-Lintuzumab downregulates the anti-apoptotic MCL-1, BCL-2 and BCL-XL protein levels in OCI-AML3 cells. OCI-AML3 cells were treated with media, unlabeled Lintuzumab or ^225^Ac-Lintuzumab (1.48 and 3.7 KBq) for 1 h, washed, and further incubated for 72 h. Cells were collected 72 h post-treatment, lysed and Western blotting was performed. (A) LI-COR Odyssey Imaging System was used to visualize the protein expression levels of MCL-1, BCL-2 and BCL-XL proteins.

### Robust anti-tumor efficacy and survival of combination therapy in venetoclax resistant xenografts

Since significant anti-tumor cell cytotoxicity was observed with combination treatment of ^225^Ac-lintuzumab and venetoclax in AML cell lines, we evaluated this combination therapy in AML subcutaneous transplantation models using OCI-AML3 and U937 xenografts in SCID mice. Vehicle-treated, venetoclax-treated and single dose of naked unlabeled lintuzumab mAbs (0.4 ug) were unable to control tumor burden in mice in both models OCI-AML3 (Fig. 5A, B, C) and U937 (Fig 6A, B, C). Single agent venetoclax treatment showed no anti-tumor activity, although a modest survival benefit was observed in these resistant models compared to vehicle control and unlabeled lintuzumab antibody. Notably, both single agent ^225^Ac-lintuzumab and ^225^Ac-lintuzumab in combination with venetoclax significantly reduced tumor burden resulting in increased survival in both OCI-AML3 (Fig. 5D, E) and U937 (Fig. 6D, E) models. Furthermore, at the end of the OCI-AML3 animal study, on day 38 post-treatment, the group treated with 7.4 KBq ^225^Ac-lintuzumab alone demonstrated 80% survival, with complete response in 2 of 5 mice (Table 1), while the group treated with the combination of ^225^Ac-lintuzumab with venetoclax showed 100% survival and CR in 3 of 5 mice (Fig. 5F). Interestingly, in the U937 animal model, there was 80% survival with the combination of 7.4 KBq ^225^Ac-lintuzumab with venetoclax and CR in 2 of 5 mice (Table 2), while the animals treated with the ^225^Ac-lintuzumab alone demonstrated 20% survival (Fig. 6F) with CR in 1 of 5 mice. Altogether these results demonstrate robust anti-tumor control and survival of the ^225^Ac-lintuzumab combination with venetoclax in AML models refractory to single agent venetoclax.

**Table 1.** Viability* and anti-tumor response^†^ to single agent and combination treatment in OCI-AML3 model.

| Treatment | D21 Response | D38 Response |
| --- | --- | --- |
| Vehicle | 0/5 Alive<br>0/5 SD<br>0/5 PR<br>0/5 CR | 0/5 Alive |
| Lintuzumab | 1/5 Alive<br>0/5 SD<br>0/5 PR<br>0/5 CR | 0/5 Alive |
| 7.4 KBq <sup>225</sup>Ac-Lintuzumab | 5/5 Alive<br>1/5 SD<br>2/5 PR<br>1/5 CR | 4/5 Alive<br>0/5 SD<br>0/5 PR<br>2/5 CR |
| Venetoclax | 4/5 Alive<br>0/5 SD<br>0/5 PR<br>0/5 CR | 0/5 Alive |
| 7.4 KBq <sup>225</sup>Ac-Lintuzumab<br>+ Venetoclax | 5/5 Alive<br>1/5 SD<br>2/5 PR<br>2/5 CR | 5/5 Alive<br>0/5 SD<br>0/5 PR<br>3/5 CR |

<sup>\*</sup> Mice were sacrificed if they experienced > 20% weight loss, tumor volume reached 4,000 mm<sup>3</sup>, or tumors became necrotic.
<sup>†</sup> Anti-tumor response was determined by change in tumor volume following treatment relative to tumor starting volume: SD, less than 30% change in tumor volume; PR, greater than 30% but less than 100% reduction in tumor volume; and CR, greater than 100% reduction in tumor volume.

**Table 2.** Viability* and anti-tumor response^†^ to single agent and combination treatment in U937 model.

| Treatment | D21 Response | D32 Response |
| --- | --- | --- |
| Vehicle | 1/5 Alive<br>0/5 SD<br>0/5 PR<br>0/5 CR | 0/5 Alive |
| Lintuzumab | 0/5 Alive<br>0/5 SD<br>0/5 PR<br>0/5 CR | 0/5 Alive |
| 7.4 KBq <sup>225</sup>Ac-Lintuzumab | 3/5 Alive<br>2/5 SD<br>0/5 PR<br>1/5 CR | 1/5 Alive<br>0/5 SD<br>0/5 PR<br>1/5 CR |
| Venetoclax | 0/5 Alive<br>0/5 SD<br>0/5 PR<br>0/5 CR | 0/5 Alive |
| 7.4 KBq <sup>225</sup>Ac-Lintuzumab+ Venetoclax | 4/5 Alive<br>1/5 SD<br>1/5 PR<br>2/5 CR | 4/5 Alive<br>0/5 SD<br>0/5 PR<br>2/5 CR |

<sup>\*</sup> Mice were sacrificed if they experienced > 20% weight loss, tumor volume reached 4,000 mm<sup>3</sup>, or tumors became necrotic.
<sup>†</sup> Anti-tumor response was determined by change in tumor volume following treatment relative to tumor starting volume: SD, less than 30% change in tumor volume; PR, greater than 30% but less than 100% reduction in tumor volume; and CR, greater than 100% reduction in tumor volume.

**Figure 5:**
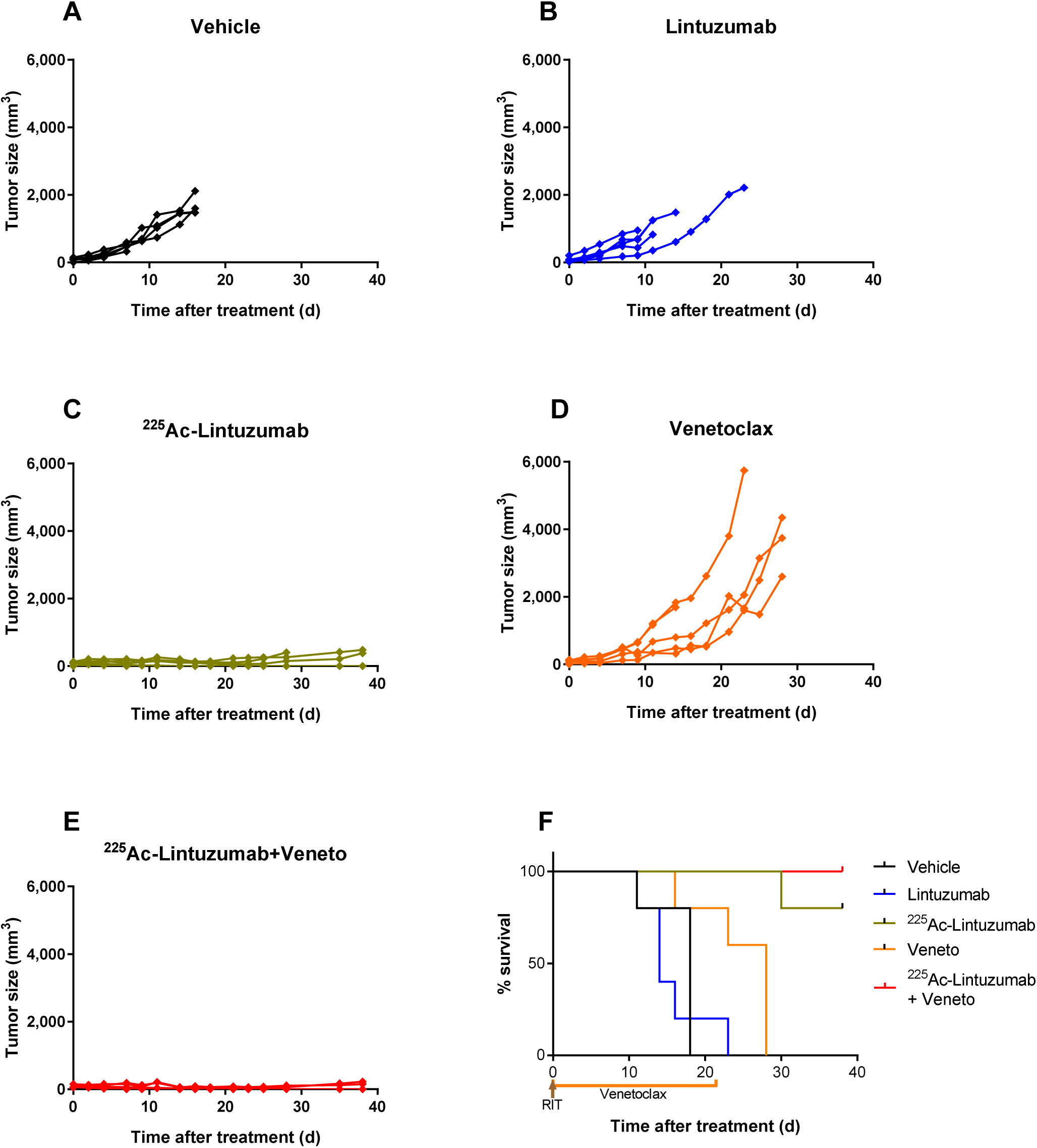
^225^Ac-Lintuzumab and venetoclax combination effects a robust anti-tumor response and increases survival benefit in OCI-AML3 xenografts. SCID mice (5 animals per group) were injected with OCI-AML3 cells into the right flank. Tumor bearing mice were treated with vehicle (A), unlabeled Lintuzumab (0.4 µg) (B), 7.4 KBq ^225^Ac-Lintuzumab (0.2 µg) (C), venetoclax (7.4 mg/kg) (D), or combination of venetoclax and 7.4 KBq ^225^Ac-Lintuzumab (E). Tumor volume was calculated using the formula V=0.5(LW^2^). (F) Kaplan-Meier graph showing animal survival.

**Figure 6:**
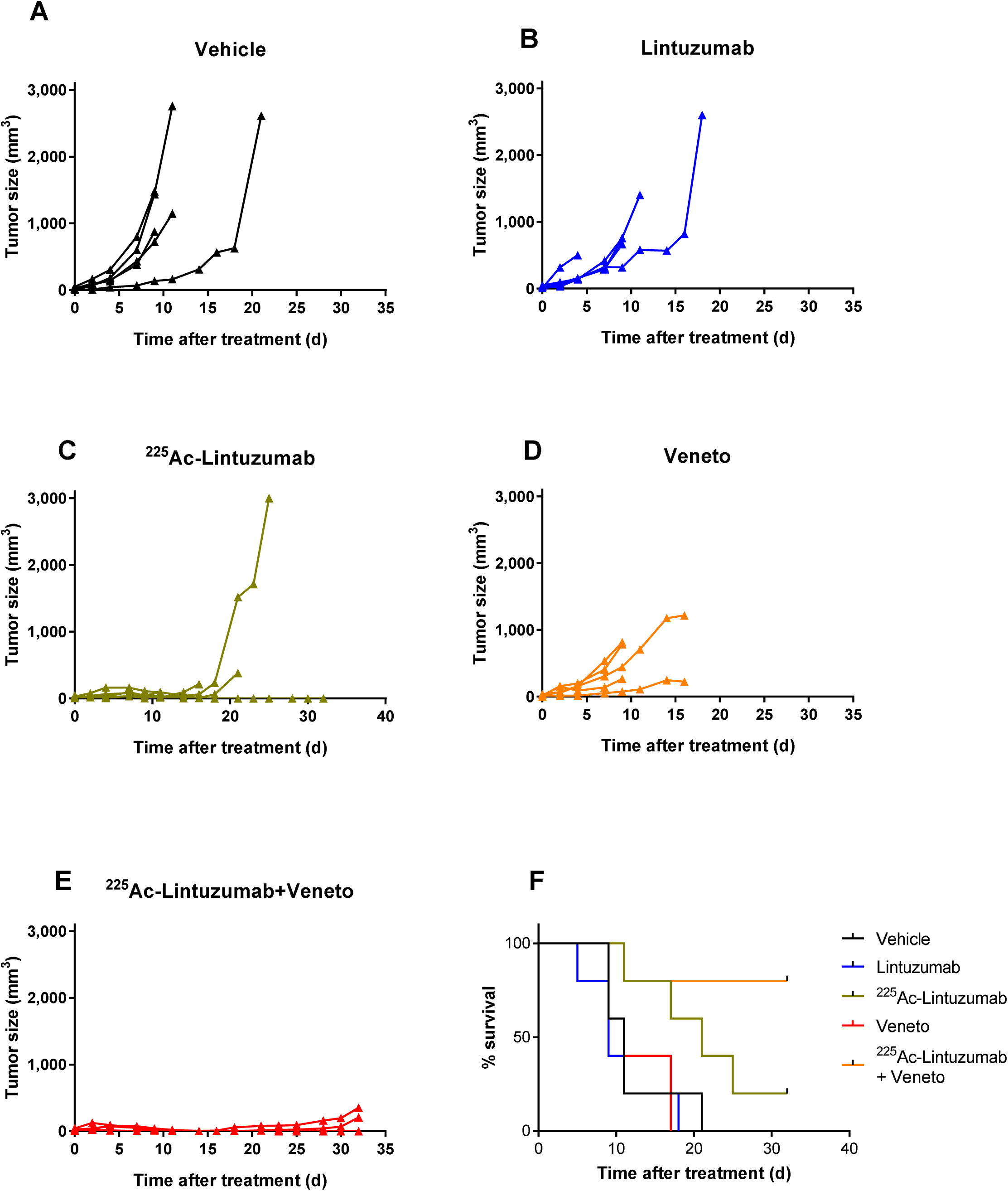
U937 xenografts model showed increased anti-tumor response and survival benefit with ^225^Ac-Lintuzumab and venetoclax combination treatment. SCID mice (5 animals per group) were injected with U937 cells into the right flank. Tumor bearing mice were treated with venetoclax (200mg/kg) (A), 7.4 KBq ^225^Ac-Lintuzumab (0.2 µg) (B), vehicle (C), unlabeled Lintuzumab (0.2 µg) (D), or combination of venetoclax and 7.4 KBq ^225^Ac-Lintuzumab (E). Tumor volume was calculated using the formula V=0.5(LW^2^). (F) Kaplan-Meier graph showing animal survival.

### Tolerable safety profile of ^225^Ac-lintuzumab and venetoclax combination in tumor-bearing mice

A safety evaluation was performed on mice treated with venetoclax or ^225^Ac-lintuzumab single agent or in combination by measuring the weight, liver toxicity (AST and ALT), kidney toxicity (Creatinine and BUN). ^225^Ac-lintuzumab alone and the combination with venetoclax demonstrated transient but reversible weight loss (Fig. 7A). There were no significant changes in AST (Fig. 7B) and ALT (Fig. 7C) levels in any of the treatment groups, though 1 of 5 mice in the venetoclax single agent cohort exhibited elevated enzyme levels. Furthermore, we did not find any evidence of increased creatinine (Fig. 7D) and BUN (Fig. 7E) levels demonstrating the absence of kidney toxicity in mice in any of the treatment groups.

**Figure 7:**
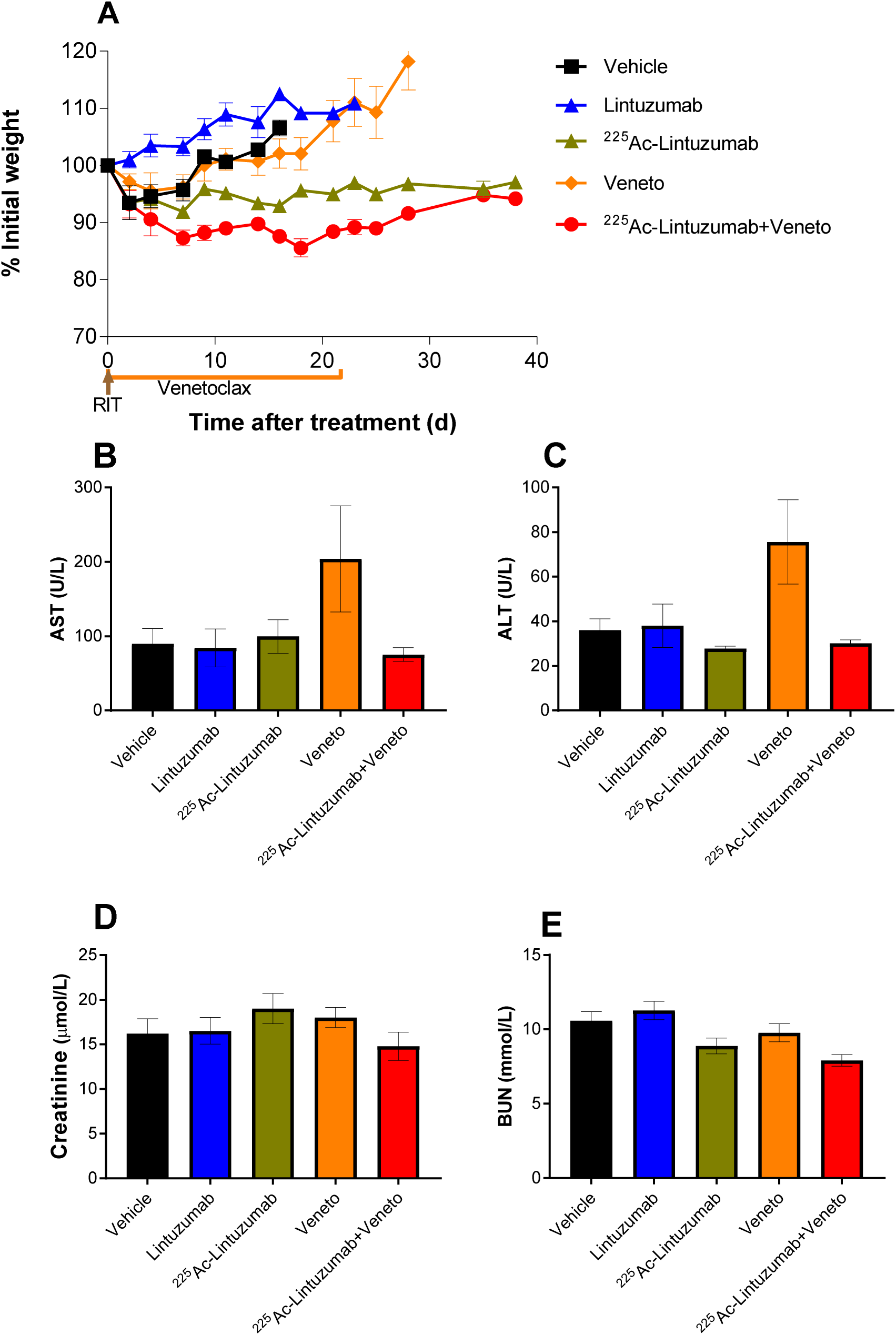
Combination treatment with ^225^Ac-Lintuzumab and venetoclax showed tolerable safety profile in tumor-bearing mice. OCI-AML3-tumor bearing mice were treated as described in the legend for Fig. 5. (A) Body weights were measured and recorded three times per week. Blood was collected from euthanized mice and analyzed for liver and kidney toxicity i.e. aspartate aminotransferase; AST (B), alanine transaminase; ALT (C), creatinine (D) and blood urea nitrogen; BUN (E). Data represents mean with SEM of 5 animals per treatment group.

## DISCUSSION

Acute myeloid leukemia (AML) is a complex hematological disease often occurring in older patients. While several new targeted therapies have been recently approved, most patients eventually relapse and succumb to their disease. Strategic clinical approaches applying rational and scientifically supported therapeutic combinations are a logical step to further extend the response to, and benefit of, these novel targeted agents. In this study, we demonstrated *in vitro* and *in vivo* that the combination of venetoclax with ^225^Ac-lintuzumab, an antibody armed with the potent alpha emitting radionuclide Actinium-225, had a synergistic effect with venetoclax on tumor cell killing in vitro and the combination provided a pronounced survival advantage in AML tumor models shown to be resistant and refractory to venetoclax.

BCL-2 family proteins can promote tumor cell survival, making them attractive drug targets ^3,26,27^. Antagonists targeting BCL-2/BCL-XL (ABT-737 and ABT-263/navitoclax) or BCL-2 only (ABT-199/GDC-0199/venetoclax) have shown clinical benefit, notably leading to the recent regulatory approval of venetoclax in CLL and AML. Not surprisingly, one mechanism of resistance to venetoclax is mediated by up-regulation or overexpression of other BCL-2 family members, including MCL-1 and BCL-XL ^7-11^. ^225^Ac-lintuzumab targets the myeloid specific marker CD33 found overexpressed on most tumor cells in AML and MDS and less frequently in MM ^17,28,29^. Phase 1 and 2 studies have demonstrated evidence of single agent clinical activity, supporting further investigation in combination with other therapies in the treatment of myeloid malignances ^18,19^. Importantly, ^225^Ac-lintuzumab internalizes on binding to its CD33 antigen into the cells and effectively carries ^225^Ac inside the cells ^22^ thereby limiting off-target toxicities and contributing to a high degree of patient tolerability in clinical trials ^17-19^. In the present study, we demonstrated robust anti-tumor activity and survival from the combination of ^225^Ac-lintuzumab with venetoclax in AML lines refractory to the BCL-2 inhibitor. It has been shown that DNA damage can reduce the expression of MCL-1 protein ^15, 30^. Therefore the use of targeted therapies that increase tumor specific DNA damage may be a promising approach to enhance the potency of venetoclax and overcome resistance mechanisms. As anticipated, exposure of AML cell lines to ^225^Ac-lintuzumab potently induced DNA double strand breaks and demonstrated single agent activity *in vitro* and *in vivo*. Importantly, the combination with venetoclax had an apparent synergistic effect on DNA damage that in turn reduced the expression of MCL-1 protein leading to a significant enhancement of anti-tumor potency in AML cell lines resistant to single agent venetoclax. To investigate the possible mechanisms by which ^225^Ac-lintuzumab may potentiate or re-sensitize resistant AML lines to the venetoclax, we assessed the impact of the antibody radio-conjugate on cellular levels of anti-apoptotic proteins MCL-1, BCL-2, and BCL-XL. Previously Niu et al.^31^ have screened eleven AML cell lines and showed wide range of venetoclax sensitivity. Based on that study, we have selected two highly venetoclax resistant (OCI-AML3 and U937) and one sensitive (MOLM-13) AML cell lines to cover the wide range of sensitivities to single-agent venetoclax – the differences between IC50 for the sensitive MOLM-13 and resistant OCI-AML3 and U937 were 131 and 78-fold, respectively. Interestingly, OCI-AML3 cells exposed to ^225^Ac-lintuzumab demonstrated significantly reduced the levels of both MCL-1 and BCL-XL compared to controls. Mechanistically, the reduction of both MCL-1 and BCL-XL was likely the result of genotoxic stress induced in tumor cells as a result of alpha particle-mediated DNA damage. The reduction of BCL-2 family proteins by ^225^Ac-lintuzumab effectively mitigates resistance to venetoclax in AML tumor lines. There might be additional mechanisms involved into the combination treatment such as radiosensitizing effect of venetoclax. In this regard, O’Steen et al. who combined radioimmunotherapy with beta-emitter ^90^Y and venetoclax, and also observed the synergistic results of such combination, refer to “multiple mechanisms” of action ^16^. The radiosensitizing nature of venetoclax warrants future studies.

As a result, the combination induced high DNA damage which in turn caused a pronounced anti-tumor response and survival benefit *in vivo* in mice bearing venetoclax resistant AML xenografts. In these models, a single dose of ^225^Ac-lintuzumab was enough to induce a durable response compared to a 21-day course of venetoclax. We anticipate that successive courses of such combination therapy would enable a sustained response, extending survival with curative intent. At present there is significant interest in the clinical evaluation of venetoclax and drug combinations with direct or indirect inhibitors of MCL-1 to overcome resistance to the BCL-2 inhibitor ^32^. Mechanistically, the combination of ^225^Ac-lintuzumab antibody radio-conjugate therapy with venetoclax offers the dual benefit of not only mediating single agent anti-tumor activity, but also, as demonstrated in this study, effects the reduction of both MCL-1 and BCL-XL. By reducing MCL-1 and BCL-XL, ^225^Ac-lintuzumab reverses resistance to venetoclax. As such, this therapeutic combination may exhibit superior response in AML in comparison with combinations solely focused on suppression of BCL-2 pathways. These results support the clinical testing of the ^225^Ac-lintuzumab in combination with venetoclax in the treatment of venetoclax-resistant AML.

## ACKNOWLEDGEMENTS

The authors would like to thank Mrs. Mackenzie Malo, Dr. Rubin Jiao and Dr. Amit Gaba for technical assistance. This study was supported by research funding from Actinium Pharmaceutical Inc. and Fedoruk Center for Nuclear Innovation.

## AUTHOR CONTRIBUTIONS

ED and DLL designed the study: RG, KJHA, WD and EG performed the experiments, RG, ED and DLL wrote the manuscript. DLL and ED came up with the concept of the study; RG, KJHA and WD performed experiments; RG, DLL, EG and ED analyzed the data; RG, DLL and ED wrote the manuscript. All authors read and approved the final manuscript.

